# Dissolvable microgel-templated macroporous hydrogels for controlled cell assembly

**DOI:** 10.1101/2022.01.05.475155

**Authors:** Zhongliang Jiang, Fang-Yi Lin, Kun Jiang, Han Nguyen, Chun-Yi Chang, Chien-Chi Lin

## Abstract

Mesenchymal stem cells (MSCs)-based therapies have been widely used to promote tissue regeneration and to modulate immune/inflammatory response. The therapeutic potential of MSCs can be further improved by forming multi-cellular spheroids. Meanwhile, hydrogels with macroporous structures are advantageous for improving mass transport properties for the cell-laden matrices. Herein, we report the fabrication of MSC-laden macroporous hydrogel scaffolds through incorporating rapidly dissolvable spherical cell-laden microgels. Dissolvable microgels were fabricated by tandem droplet-microfluidics and thiol-norbornene photopolymerization using a novel fast-degrading macromer poly(ethylene glycol)-norbornene-dopamine (PEGNB-Dopa). The cell-laden microgels were subsequently encapsulated within another bulk hydrogel matrix, whose porous structure was generated efficiently by the rapid degradation of the PEGNB-Dopa microgels. The cytocompatibility of this *in situ* pore-forming approach was demonstrated with multiple cell types. Furthermore, adjusting the stiffness and cell adhesiveness of the bulk hydrogels afforded the formation of solid cell spheroids or hollow spheres. The assembly of solid or hollow MSC spheroids led to differential activation of AKT pathway. Finally, MSCs solid spheroids formed *in situ* within the macroporous hydrogels exhibited robust secretion of HGF, VEGF-A, IL-6, IL-8, and TIMP-2. In summary, this platform provides an innovative method for forming cell-laden macroporous hydrogels for a variety of future biomedical applications.

## 1. Introduction

Multipotent mesenchymal stem cells (MSCs) have high potential for many cell-based therapies as they can be isolated from multiple sources (e.g., bone marrow, umbilical cord, adipose tissue, etc.) [1] and be readily differentiated into adipocytes, osteoblasts, and chondrocytes [2]. MSCs can also undergo transdifferentiation into cardiomyocytes [3], neurons [4], and corneal epithelium cells [5]. Additionally, studies have found that the secretory factors of MSCs can be utilized for regulating inflammatory and immune responses [6, 7]. Notably, MSC secretome can be manipulated by controlling marix stiffness, viscoelasticity, and the presence of biological motifs (e.g., fibronectin) [8-12]. Many MSC-secreted factors, including transforming growth factor-β (TGF-β), insulin-like growth factor (IGF), and fibroblast growth factor-2 (FGF-2), have been shown to stimulate cell proliferation, migration, and differentiation [12-15]. Furthermore, studies have found that, compared with 2D culture, MSCs cultured in 3D spheroids promoted cell proliferation, secretory properties, cell-cell interactions, and *in vivo* survival [16]. In particular, a 2D surface un-naturally polarizes cells and lacks characteristics/architectures of a native cellular microenvironment needed to maintain the phenotype of the isolated cells [16]. Studies have also revealed that assembling MSCs into multi-cellular spheroids increased the level of cell-cell interaction, leading to dramatically improved secretory properties and differentiation potential towards certain lineages (e.g., hepatocytes [17], osteoblasts [18], and neurons [19]). MSCs have also been reported to exhibit anti-cancer effects via enhancing AKT signaling [20].

3D hydrogel scaffolds with physicochemically relevant properties have increasingly being used for MSC culture [21-23]. Unless using hydrogel amenable to controllable degradation, chemically crosslinked hydrogels generally have nanometer-scale pores (or mesh) that will impede macromolecular diffusivity [24, 25]. Additionally, MSCs do not naturally form spheroids in 3D hydrogels, hence the encapsulated MSCs would remain disperse within the hydrogel matrix. The most efficient method to generate MSC clusters is by using well-plates with a hydrophobic surface and/or with special geometry (e.g., AggreWell™ plate) [26]. After assembly, MSC spheroids are collected and encapsulated in a hydrogel matrix to provide the cells with desired cell-matrix interactions. However, the handling of MSC spheroids may be challenging and the distribution of MSC spheroids within the bulk hydrogels may be non-uniform due to the settlement of the multicell spheroids. Furthermore, very few methods permit the generation of hollow cell spheres with a lumen structure, such as that found in ocular lens and lung alveoli [27]. Therefore, a robust and powerful material engineering platform is needed to assist structured MSCs assembly *in situ*.

Hydrogels with macroscopic pores (i.e., macroporous hydrogels) are ideal for many biomedical applications owing to their improved transport properties [28] and opportunity for enhanced cell-cell interactions within the matrix [29]. Given the biphasic nature of the macroporous hydrogels, extracellular properties of the encapsulated cells could be decoupled from the bulk hydrogel matrix characteristics. As such, cellular assembly, attachment, and differentiation could be controlled by tuning extracellular properties independent of hydrogel stiffness, viscoelasticity, and cell adhesiveness [30-32]. A number of approaches have been employed to fabricate macroporous hydrogels, including salt leaching and gas forming, which depend on embedding and removing porogens from the crosslinked hydrogels [33]. While this approach is straightforward, commonly used porogens (e.g., NaCl crystals and CO_2_) are not compatible with *in situ* cell encapsulation. Furthermore, post-synthesis cell seeding is required but the passive penetration of cells into the porous hydrogel may be challenging, resulting in non-uniform cell distribution. Alternatively, cell-laden microgel templating within a continuous matrix could directly introduce cells into a 3D environment and cell structures could be formed *in situ*. But sacrificial materials (e.g., pH, photo, enzyme or thermal sensitive) are either limited in cytocompatibility or their degradation products (e.g., gelatin-derived peptides) may alter stem cell signaling pathways [34, 35]. Even though gelatin and alginate based hydrogels/microgels offer good cytocompatibility to the encapsulated cells, their dissolution requires changing temperature (for gelatin) or using external molecular trigger (e.g., metal ion chelators).

Recently, microporous annealed particle hydrogels (MAP) have become an attractive strategy where MSCs were seeded and attached on the annealed microgel interfaces [36]. However, current MAP scaffolds cannot support MSC assembly into spheroids. Alternatively, hydrogels susceptible to hydrolytic degradation could be harnessed for forming dissolvable microgels to template the otherwise nanoporous hydrogels into macroporous gels. For example, microgels crosslinked from poly(ethylene glycol) (PEG) diacrylate (PEGDA) and dithiol crosslinker have been templated as sacrificial phase. The ester bonds in crosslinked microgels are liable for hydrolysis for fabricating macroporous hydrogels [37]. However, conventional ester hydrolysis induced gel degradation is a slow process that precludes the applications of such platform in sensitive cell candidates that rely on cell-cell interaction to survive and function.

Our group has recently developed a new class of PEG-based macromers (e.g., PEG-norbornene-dopamin or PEGNB-Dopa) crosslinkable by light induced thiol-norbornene crosslinking and degradable via ultra-fast hydrolytic degradation kinetics [38]. For instance, photopolymerized PEGNB-Dopa/dithiothreitol (DTT) hydrogels with an initial stiffness as high as 10 kPa were degraded completely within 2 hours upon contact with buffer solution. In this study, we employed droplet-microfluidics to fabricate dissolvable PEGNB-Dopa microgels with well-controlled size and degradation profiles. These cell-laden dissolvable microgels were loaded into another bulk hydrogel. Macroporous hydrogels were formed after the dissolution of the microgels, leaving behind the cells in the cavities. By tuning cell adhesiveness of the continuous matrix, multi-cellular structures were formed with pancreatic cancer cells COLO-357, lung epithelial cells A549, as well as mouse and human MSCs. Besides morphological tunability, we also investigated whether different assembly structures affect MSC therapeutic potential by analyzing phosphorylation of a panel of AKT signaling proteins, as well as secretion of growth factors and cytokines.

## 2. Results and Discussion

### 2.1. Degradation of PEGNB-Dopa hydrogels

PEGNB-Dopa, a new derivative of PEGNB macromer recently developed in our group, was chosen for forming dissolvable microgels.[38] PEGNB-Dopa possesses the same rapid and orthogonal crosslinking efficiency afforded by the cytocompatible thiol-norbornene photopolymerization [39, 40]. Once crosslinked, PEGNB-Dopa based hydrogels would degrade rapidly in aqueous buffer at 37°C via enhanced ester hydrolysis (**Figure 1A**). Using PEG4SH as the crosslinker at different thiol/ene ratio, we first evaluated the photocrosslinking of PEGNB-Dopa into thiol-norbornene hydrogels. As with other thiol-norbornene hydrogels, the degree of hydrogel crosslinking scales with the thiol/ene ratio, with the highest shear moduli (G’) reaching 16 kPa at a stoichiometric ratio (**Figure 1B**). Regardless of initial crosslinking density (i.e., G’ ∼ 1.2 kPa to 16 kPa), all PEGNB-Dopa crosslinked hydrogels degraded rapidly in aqueous buffer, with gels crosslinked at low thiol/ene ratios degraded faster than at high thiol/ene ratios (**Figure 1C**). For example, at a thiol/ene ratio of 0.5, PEGNB-Dopa/PEG4SH hydrogels degraded completely within 2 hours, a result similar to our prior report.[38] In contrast, hydrogels crosslinked by conventional PEGNB did not exhibit any noticeable degradation within 8 hours. It is important to note that even hydrogels with high initial modulus (G’ > 16 kPa) could be completely degraded within 6 hours after incubating in water (**Figure 1C**). We have previously reported the accelerated ester hydrolysis kinetics of PEGNB-Dopa hydrogels, with a pseudo-first order degradation kinetic constant (k_hyd_) of less than 1 hr^-1^. This is in stark contrast to the conventional hydrolytically degradable thiol-norbornene hydrogels that could only be degraded hydolytically over weeks or even months [41]. Given that stiffer hydrogels can be easily handled and processed in a microfluidic droplet generator, PEGNB-Dopa hydrogels crosslinked at stoichiometric thiol/ene ratio were used for the following experiments.

**Figure 1.**
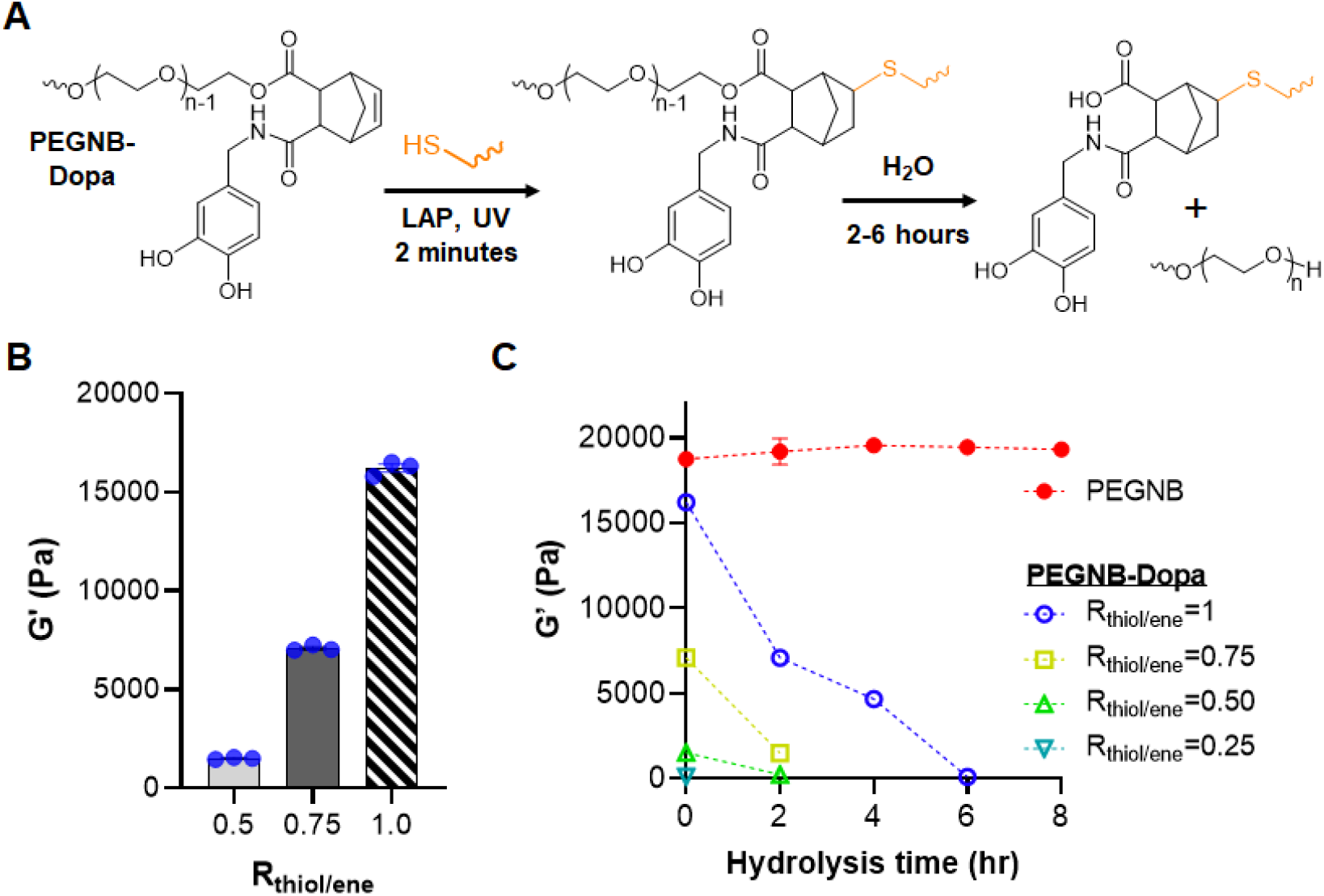
Tunable crosslinking and rapid degradation of PEGNB-Dopa hydrogels. (A) Chemical structure of PEGNB-Dopa along with the schematic of its gelation and hydrolytic degradation. (B) Initial shear moduli of PEGNB-Dopa hydrogels formed at different thiol/ene ratio. (C) Degradation of PEGNB and PEGNB-Dopa hydrogels formed with different thiol/ene ratios as indicated by the decrease of shear modulus over time (n=3, Mean ± SEM).

### 2.2. Characterization of macroporous hydrogels

We reasoned that the rapid degradation of PEGNB-Dopa hydrogels could be formulated into dissolvable microgels, serving as sacrificial templates for forming macroporous hydrogels. To achieve this, we employed tandem droplet-microfluidics and thiol-norbornene photopolymerization to synthesize near-monodispersed PEGNB-Dopa microgels (**Figure 2A**). Next, the PEGNB-Dopa microgels (crosslinked in the presence of thiolated Rhodamine B for visualization) [42] were collected and further encapsulated in another bulk hydrogel (**Figure 2B**). Of note, the bulk hydrogels were crosslinked by the same thiol-norbornene photopolymerization using conventional PEGNB, followed by incubating the composite gels in water to allow for dissolution of the encapsulated microgels. To facilitate the visualization of macro-scale pore formation, 0.1 wt% FITC-PEG-SH was added during the bulk hydrogel crosslinking. Confocal imaging revealed that the rhodamine-labeled microgels were dispersed in the FITC-labeled bulk gel (**Figure 2C**, top panel). After 24 hours of incubation in PBS at 37°C, minimal red fluorescence was detected, indicating the successful dissolution of the microgels (**Figure 2C**, bottom panel). As such, macroporous hydrogels were obtained without the use of any solvent or change in temperature for the removal of the microgel porogens. It can be seen from the confocal images that some microgels were in close contact, resulting in some level of interconnective of the pores. Future work may explore jam-packing the microgels within the bulk gels to create highly interconnected macroporous scaffold. Control experiment using non-degradable PEGNB microgels resulted in minimum change of fluorescence even after 168 hours of incubation (**Figure S1**). The formation of cavities with PEGNB-Dopa microgels was further confirmed by SEM imaging, where large pores were only observed in PEGNB-Dopa microgel templated hydrogels (**Figure 2D, 2E**). Tracking moduli of the composite hydrogels revealed that the degradation process was completed within 24 hours, as stable moduli were achieved afterwards (**Figure S2**). Stiffness of the macroporous hydrogels was readily tuned depending on continuous matrix employed, in this case, PEGNB or gelatin-norbornene (GelNB) with 0.5 or 1 thiol/ene ratio (**Figure S2A**). Increasing the volume fraction of PEGNB-Dopa microgels within the continuous matrix (0%, 50%, or 68%) led to a slight increase in swelling ratio (**Figure S2B**), indicating that the creation of the pores did not significantly alter the macroscopic hydrogel mechanical properties. Taken together, these results indicate success fabrication of macroporous hydrogels through using dissolvable PEGNB-Dopa microgels.

**Figure 2.**
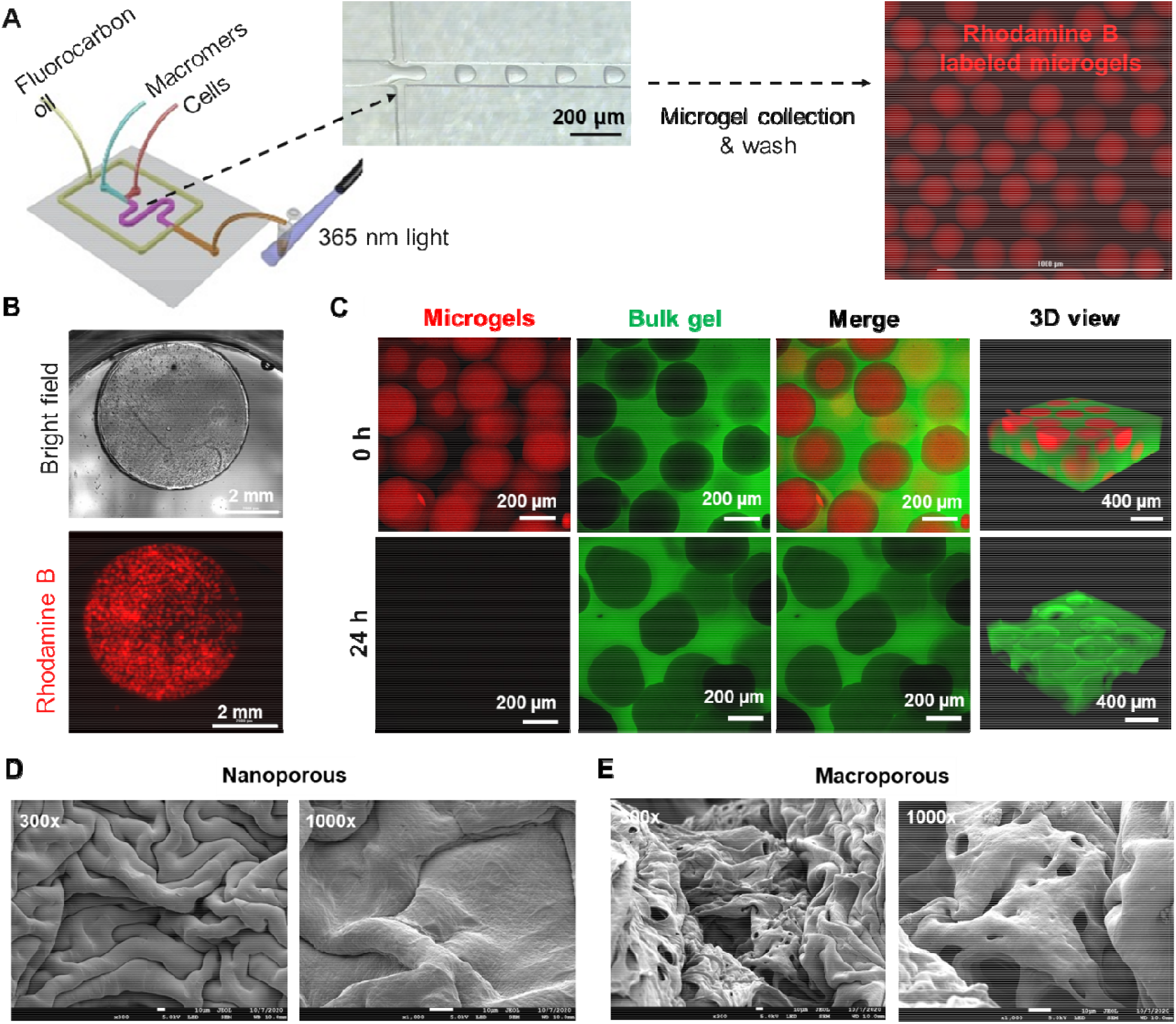
Fabrication and characterization of macroporous hydrogels. (A) Schematic of tandem microfluidic droplet generation and thiol-norbornene photopolymerization to form monodisperse microgels. (B) Bright field and fluorescence images of a microgel-templated hydrogel. (C) Confocal images of the microgel-templated hydrogel. The dissolution of the microgels resulted in a macroporous hydrogel. (D, E) SEM images of nanoporous (D) and macroporous (E) hydrogels. Scale bar = 10 μm.

### 2.3. In situ formation of tumor spheorids in macroporous hydrogels

Droplet microfluidic devices have been used for *in situ* encapsulation of cells [43-45] and were employed to encapsulate cells within the PEGNB-Dopa microgels. An advantage of using droplet microfluidic device to fabricate microgels is that the size of the microgels can be precisely controlled by adjusting the flow rates of the oil and/or aqueous phases. For example, at a constant aqueous phase flow rate of 4 μl/min, decreasing oil phase flow rate from 20, 12, to 4 μl/min led to increasing diameter of the microgels (ca. 150 μm, 170 μm and 200 μm, respectively) (**Figure 3A, 3C**). We demonstrated the cytocompatibility of this process by *in situ* encapsulation of A549 lung epithelial cells in the microgels, followed by a secondary encapsulation of the cell-laden microgels in a non-degradable, bio-inert PEGNB hydrogel (**Figure 3B**). Live/dead staining results showed that A549 cells entrapped in the macroporous hydrogels cells remained alive and formed loosely packed cell clusters within the porous bulk gel day 1 post-encapsulation (**Figure 3B**). Since non-degradable, bio-inert PEGNB was employed as the continuous matrix, the entrapped cells were not provided with an adhesive surface and were forced into spheroids at concave interfaces as microgels degraded. Over 21 days of culture, the cells assembled and grew into larger solid spheroids (**Figure 3B, S3A**). When multiple microgels coalesced together, irregularly shaped cavities formed following the dissolution of the microgels. As a result, spheroids formed with irregular and elongated shape (**Figure 3B**, arrow). Note that the spheroid size is dependent on the size of cavities, which is ultimately controlled by the size of the dissolvable microgels (**Figure 3C**). Increasing microgel diameters led to enlarged cavities and therefore, larger spheroids. The process was compatible with other cell types, including a pancreatic cancer cell line, COLO-357 (**Figure S3B, S3C**).

**Figure 3.**
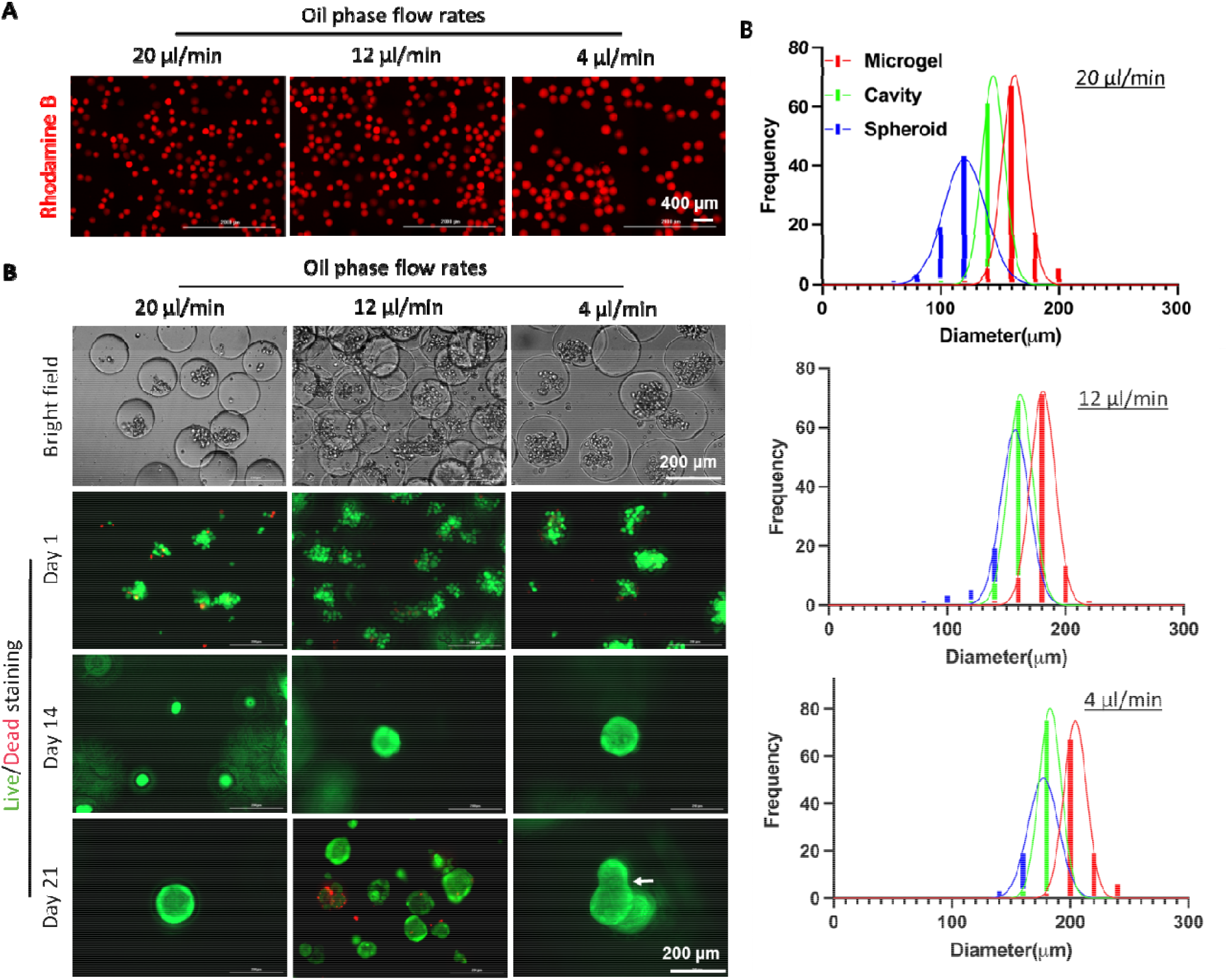
A549 spheroid formation within macroporous hydrogels. (A) Microgels with different sizes were synthesized by controlling flow rate of the oil phase while keeping flow rate of the macromer phase at 4 μl/min. (B) Bright field images (day 1) and Live/Dead staining of A549 cells on day 1, 14, and 21 post-encapsulation. (C) Quantification of the diameter of microgels, formed cavities, and spheroids as a function of oil phase flow rate.

### 2.4. Assembly of solid spheroids from mMSCs using PEGNB as continuous matrix

Emerging evidence has shown that the secretome of MSC spheroids differs drastically from that of single cells [16, 46]. However, unlike cancer cells that prefer growing into clusters in 3D hydrogels [47, 48]. MSCs do not naturally grow into spheroids in chemically crosslinked gels [49], necessitating an engineering strategy for aggregating the cells. By encapsulating MSCs into our macroporous hydrogels, we assumed MSCs would undergo a self-assembly process similarly as the A549 cells did. To prevent settlement of cells in the syringes during emulsification, density of the aqueous phase containing cells was adjusted (with OptiPrep, Sigma Aldrich) to match that of the cells (ca. 1.06 mg/mL). Failure to minimize fluid density differences would result in cells either settling down or aggregating in the syringe, reducing uniformity of the cell-to-droplet ratio. The ability to maintain cell viability while dispersing them within the carrying media for hours is crucial for scaling up the production of cell-laden microgels. Unlike cancer cells that tend to grow as spheroids, the ultimate size of engineered MSC spheroids depends on the initial number of cells used instead of microgel size.

We first utilized mouse MSC (mMSC) to demonstrate *in situ* formation of MSC spheroids in the macroporous hydrogels. mMSC density was adjusted from 2×10^6^ cells/mL to 2×10^7^ cells/mL. The flow rates for cell stream, macromer stream, and oil stream were kept constant at 2 μl/min, 2 μl/min, and 20 μl/min, respectively. Above 90% of the encapsulated cells remained alive following the microfluidic encapsulation process (**Figure 4A**). The microgel-encapsulated mMSCs formed spheroids over 7 days of culture within a continuous matrix crosslinked by non-degradable PEGNB and PEG4SH. The majority of the cells remained alive in the macroporous hydrogels, indicating the cytocompatibility of this platform to support the formation of solid and viable MSC spheroids (**Figure 4A**). Cytoskeleton staining of F-actin and nuclei revealed that assembled MSC clusters have spherical morphology with limited spreading/invasion into the bulk matrix (**Figure 4A**). Additionally, mMSC spheroids sizes were positively correlated to the cell density. Specifically, increasing cell concentration from 2×10^6^ cells/mL to 2×10^7^ cells/mL led to MSC spheroids averaging 20 to 80 μm in diameter (**Figure 4B**). Furthermore, we found that the sizes of mMSC spheroids were all smaller than the sizes of the cavities/pores, which themselves were slightly smaller than the sizes of the microgels (**Figure 4C**). The smaller cavity/pore size was a result of bulk hydrogel swelling, whereas the much smaller spheroids may be attributed to the limited proliferation of mMSC spheroids. To further enlarge the size of MSC spheroids, cells may be further concentrated within the carrying media. Alternatively, microgel size could be reduced to increase the cell-to-droplet ratio. One of the other benefits for using this platform to form macroporous hydrogels is that the mechanical or physiological properties of the continuous matrix can be easily customized to induce differential cell behaviors, such as cell attachment and differentiation. When thiol/ene ratio in continuous non-degradable PEGNB hydrogel was lowered to 0.5, overall stiffness of the hydrogels was decreased to 5 kPa (**Figure, S2A**). However, MSC spheroids were formed identically, in terms of size and shape, to those formed in the stiffer bulk hydrogels (data not shown), presumably due to limited cell-matrix interactions as the cavity sizes were considerably larger than the spheroid sizes (**Figure 4B, 4C**).

**Figure 4.**
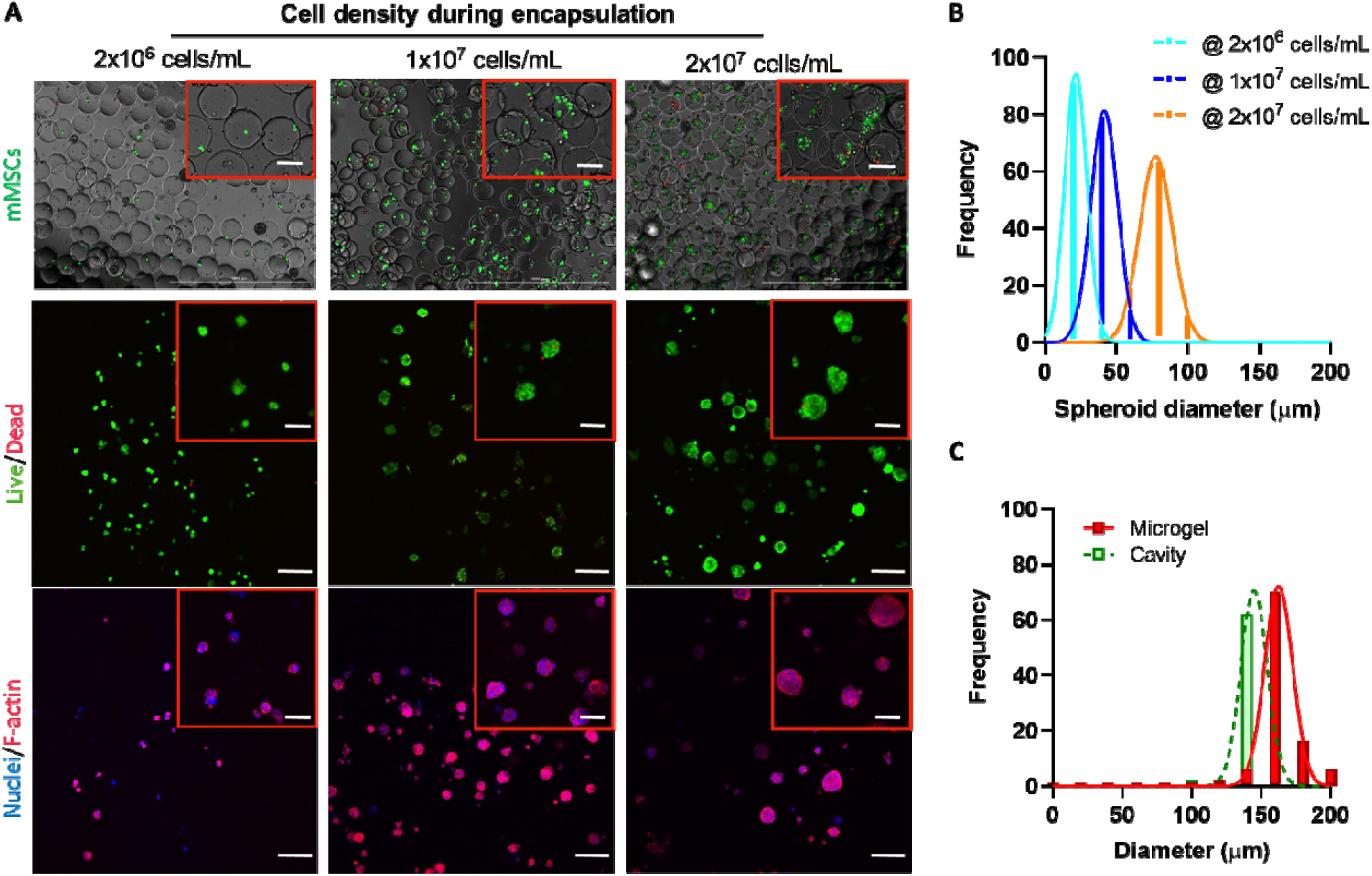
mMSC solid spheroid formation within macroporous hydrogels. (A) Live/Dead staining of mMSCs immediately after microencapsulation into PEGNB-Dopa microgels. mMSCs with a density of 2×10^6^ cells/mL, 1×10^7^ cells/mL, and 2×10^7^ cells/mL in OptiPrep were used for encapsulation. (B) Live/Dead and cytoskeleton fluorescence staining of mMSC spheroids after 7 days of culture. Scale bar = 100 μm. (C) Quantification of the sizes of microgels, cavities, and mMSC spheroids formed with various cell loading concentrations after 7 days of culture.

### 2.5. Assembly of hollow spheres from mMSCs using GelNB as continuous matrix

When a non-degradable inert PEGNB hydrogel was used as the continuous matrix, there was limited cell-matrix interaction, which facilitated solid multicellular cluster formation (**Figure 4**). When cell adhesive and protease-degradable GelNB [49] was used to encapsulate MSC-laden microgels, cells were allowed to attach and degrade the bulk matrix following the dissolution of the PEGNB-Dopa microgels. Indeed, after 7 days of incubation, MSCs stayed alive and appeared attaching to the continuous GelNB matrix and formed hollow spheres containing a lumen, as confirmed by confocal imaging (**Figure S4A**). We believe that cell migration was not an important factor in the initial phase of lumen formation owing to rapid degradation of the bioinert PEGNB-Dopa microgels (**Figure 1C**). Instead, the encapsulated cells were ‘liberated’ from the degrading microgels and simply ‘fell’ onto the surface of the continuous matrix. The ‘liberated’ cells were allowed to adhere to the surface of the cell-adhesive GelNB hydrogel. Notably, cellular protrusion into the continuous matrix was observed (**Figure S4A**, arrow), indicating mMSCs can degrade their local network to a certain degree that allows cell penetration into the matrix. Interestingly, the ability of the encapsulated cells to form hollow cell spheres was affected by the shear modulus of the bulk hydrogels as lowering the bulk gel modulus (R = 0.5) led to extensive cell spreading and the formation of an interconnected cellular network (**Figure S4B**). This was because the soft bulk GelNB hydrogels permitted MSCs to rapidly degrade and invade the surrounding matrix. In contrast, MSCs encapsulated in nanoporous PEGNB hydrogels exhibited a round morphology, while they had a more spreading morphology in GelNB hydrogels (**Figure S5A**). The results indicate that within our macroporous hydrogels, the competition of cell-cell and cell-matrix interactions could be controlled by tuning bulk matrix adhesiveness and stiffness. It is worth noting that in this contribution we only utilized bioinert (PEGNB) and bioactive (GelNB) macromers for the crosslinking of bulk hydrogels. However, the unique ability to customize cell assembly structures using degradaing PEGNB-Dopa microgels could make this platform an enabling technique for other biomaterials (e.g., alginate or RGD-conjugated alginate), which may find extensive applications in engineering lumen-containing tissues in a geometrically defined environment.

### 2.6. Evaluation of cell structures formed with mouse or human MSCs

To investigate whether cellular structures would affect cell phenotype, E-cadherin and N-cadherin were stained in MSC solid spheroids and hollow spheres. Both E- and N-cadherin were detected in MSC solid spheroids and hollow spheres (**Figure 5A**), indicating these structured MSCs still preserved both epithelial and mesenchymal phenotypes. The simultaneous expression of both cadherins indicates the quiescent state of the structured MSCs [50, 51], and they may be favorably induced into functional cell types following differentiation or transdifferentiation. A single confocal image at the middle section revealed distinct cellular structures with either solid or hollow interior (**Figure 5A**).

**Figure 5.**
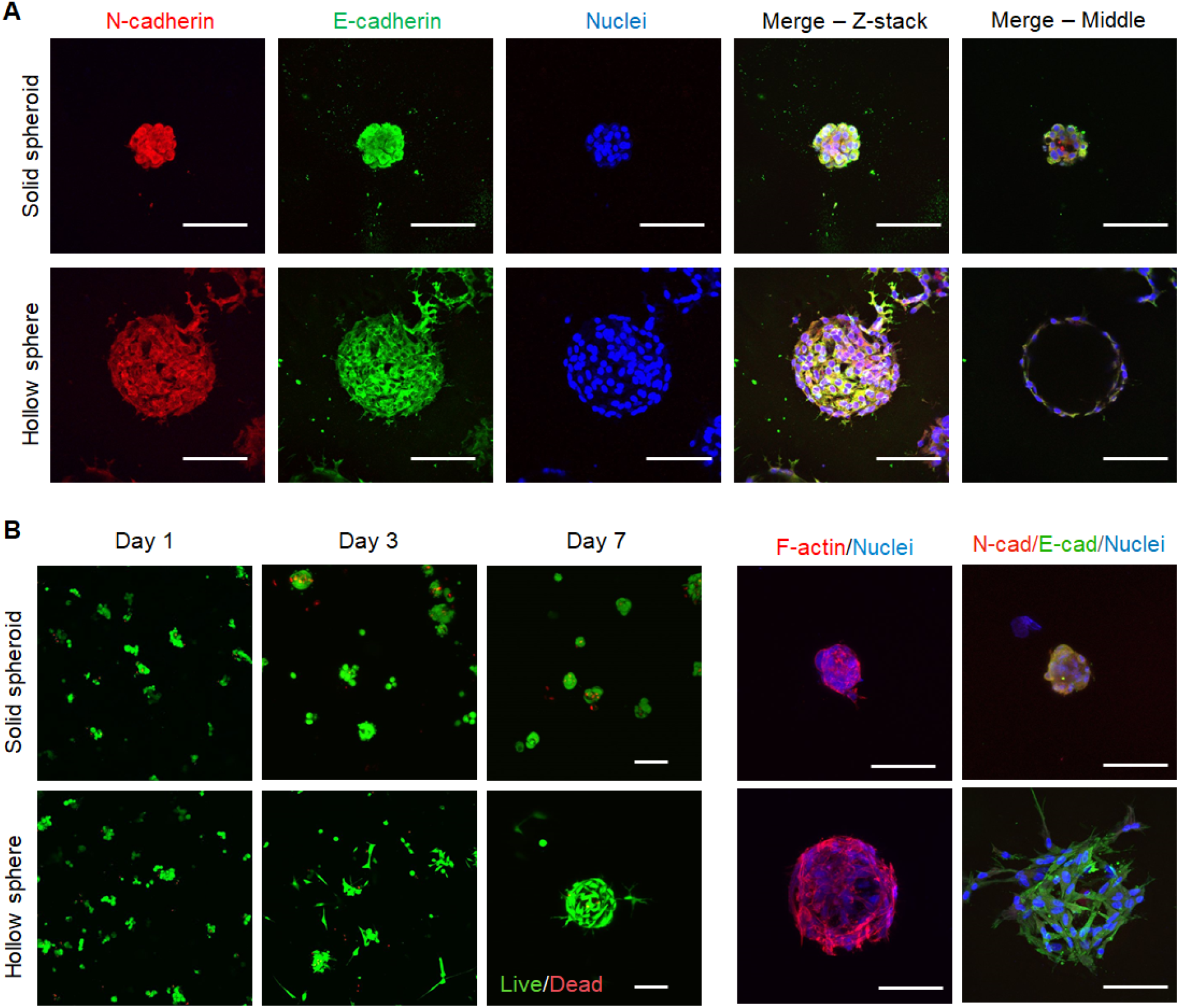
Fluorescence imaging of solid spheroids and hollow spheres formed with MSCs. (A) Immunofluorescence staining of E/N-cadherin and nuclei in mMSC solid spheroids or hollow spheres on day 7. (B) Live/Dead staining and immunofluorescence staining of cytoskeleton, E/N-cadherin and nuclei in hMSC solid spheroids or hollow spheres. Scale bar = 100 μm.

We also used human MSCs (hMSCs) to investigate whether this cell-assembly platform could be applied to human cells, which were typically larger than murine cells. We found that hMSCs also survived the macroporous hydrogel forming process with a viability above 90% in both cases (**Figure S5B**) and self-assembled into either solid spheroids or hollow spheres, depending on the cell adhesiveness of the continuous matrix (**Figure 5B**). It is worth noting that hMSCs did not proliferate when forming spheorids, while they can proliferate in the form of hollow spheres, as indicated by fluorescent images (**Figure 5B**) and DNA quantification (**Figure S5C**). Meanwhile, structured hMSCs also preserve epithelial and mesenchymal phenotypes, as indicated by the expression of E- and N-cadherin (**Figure 5B**). Importantly, both cadherins have been related with MSC paracrine activity and fate decision, thus, the strong expression of E/N-cadherin in structured MSCs may affect their therapeutic potentials [52, 53].

### 2.7. AKT pathway activation in hMSCs with different 3D growth pattern

MSCs can sense and response to the properties of the extracellular matrix via activating intracellular signaling pathways [22, 23]. Additionally, recent studies have discovered that MSC spheroids exert drastically different trophic factor secretion profiles due to heightened cell-cell interactions [21]. Therefore, structured/assembled MSCs could be highly advantageous toward engineering MSC-based therapeutics. To this end, we extracted cell lysates from hMSCs cultured in spheroids and hollow spheres formed in macroporous hydrogels and evaluated phosphorylation levels of 19 proteins involved in the AKT signaling pathway. hMSCs cultured on 2D TCPs and encapsulated in 3D nanoporous hydrogels were used as controls (**Figure 6A**). AKT pathway was studied here due to it’s diverse role in cell proliferation, survival, and apoptosis. We analyzed the levels of protein phosphorylation and grouped them into two heat maps with high (**Figure 6B**) and low (**Figure 6C**) signals for easy visualization the differences between proteins and culture conditions. Overall, the phosphorylation levels of the AKT pathway proteins in hMSC hollow spheres were similar to that in 2D culture, while the AKT protein phosphorylation levels in 3D single cells were similar to that in spheroids. Another phenomenon worth noting was the high level of phosphorylation of many AKT pathway proteins in 3D hMSC cultures, both in solid spheroids and direct encapsulation of dispersed single cells. While endogenous (i.e., without using an activator) phosphorylation of AKT was similar among the four groups (**Figure 6B**), downstream proteins that are positively regulated by AKT phosphorylation, including mTOR, P70S6K, and RPS6, had higher phosphorylation in single cells and solid spheroids than in 2D and hollow spheres. In addition, the phosphorylation of PDK1, a growth-promoting protein upstream of AKT activation [54], was similar between solid spheroids and single cells, but substaintially lower in hollow spheres. On the other hand, the phosphorylation of PTEN, a growth-inhibiting protein upstream of AKT [55], was highest in the single cells group, indicating that the lack of cell-cell interaction is unfavorable for proliferation of hMSCs in 3D. Another notable difference was the higher phosphorylation of p27 and P53 in single cells and solid spheroids. p27 inhibits cell cycle progression, while P53 is a known growth suppressor. The higher phosphorylation of these two proteins in 3D single hMSCs and spheroids suggested that the cells in these formats may experience lower proliferation capacity. Thess results suggest that both cell-materials and cell-cell interactions affect the autonomous activation of AKT signaling. Of note, both hMSCs cultured in 2D TCP and in hollow spheres in macroporous hydrogels formed monolayer-like structures with poralized cell-cell interactions (i.e., an apical surface facing media/cavity and a basal surface adhering to TCP/hydrogel). On the other hand, hMSCs encapsulated as single cells or assembled in spheroids did not exhibit substantial epical/basal polarization (**Figure 5**). The apical/basal polarity is implicated in stem cell fate processes. Future studies may focus on exploring the relationship between AKT pathway activation on hMSC proliferation and differentiation in various forms of 3D structures.

**Figure 6.**
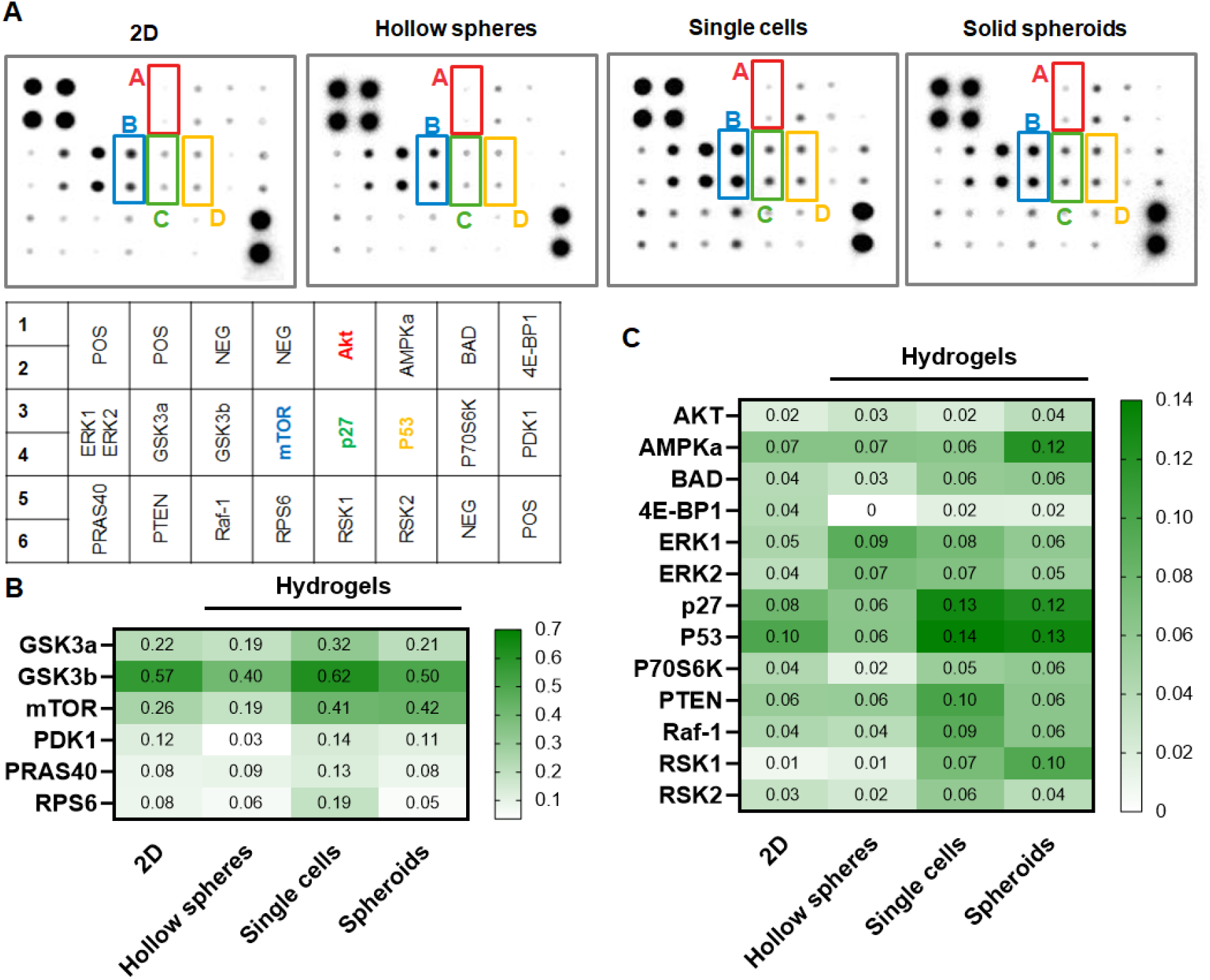
AKT phosphorylation array. (A) Blotted membranes using hMSC lysates from 2D TCPs, 3D encapsulated single cells, as well as hollow spheres or solid spheroids formed in macroporous hydrogels. (B, C) Heatmaps of phosphorylation level of AKT pathway.

### 2.8. Secretory properties of assembeled hMSCs

To further evaluate whether assembly of hMSCs into solid spheroids would improve secretory properties, we collected conditioned media (CM) from hMSCs encapsulated as single cells or formed solid spheroids in the macroporous hydrogels. These two conditions were selected for comparison for their similarity in AKT pathway activation (**Figure 6**). CM were subjected to antibody array for detecting arrays of growth factors and inflamatory cytokines (**Figure S6**). We found that the secretion of several growth factors and receptors, including FGF-6, FGF-7, HGF, IGFBP-1, IGFBP-4, IGFBP-6, IGF-1R, PLGF, SCF, TGFβ2, and VEGF-A were noticeably enhanced in hMSC spheroids than in direct 3D cell encapsulation (**Figure 7A**). Additionally, hMSCs also secreted higher amount of IL-6, IL-8, and TIMP-2 once they were assembled into spheroids (**Figure 7B**). IL-6 and IL-8 play critical roles in regulating inflammatory response [56], reaffirming that hMSC spheroids can be leveraged to regulate regeneration via their secretomes. On the other hand, the increased secretion of TIMP-2, which inhibited activity of matrix metalloproteinase 2 (MMP2) [57], suggested that the assembled hMSC speroids exhibited reduced migratory potential due to a reduced level of MMP activity. While this panels of growth factor and inflammation cytokine arrays only represent a fraction of MSC secretomes, it is reasonable to suggest, based on the current results, that the hMSC spheroids secretomes could be beneficial for regenerative medicine applications.

**Figure 7.**
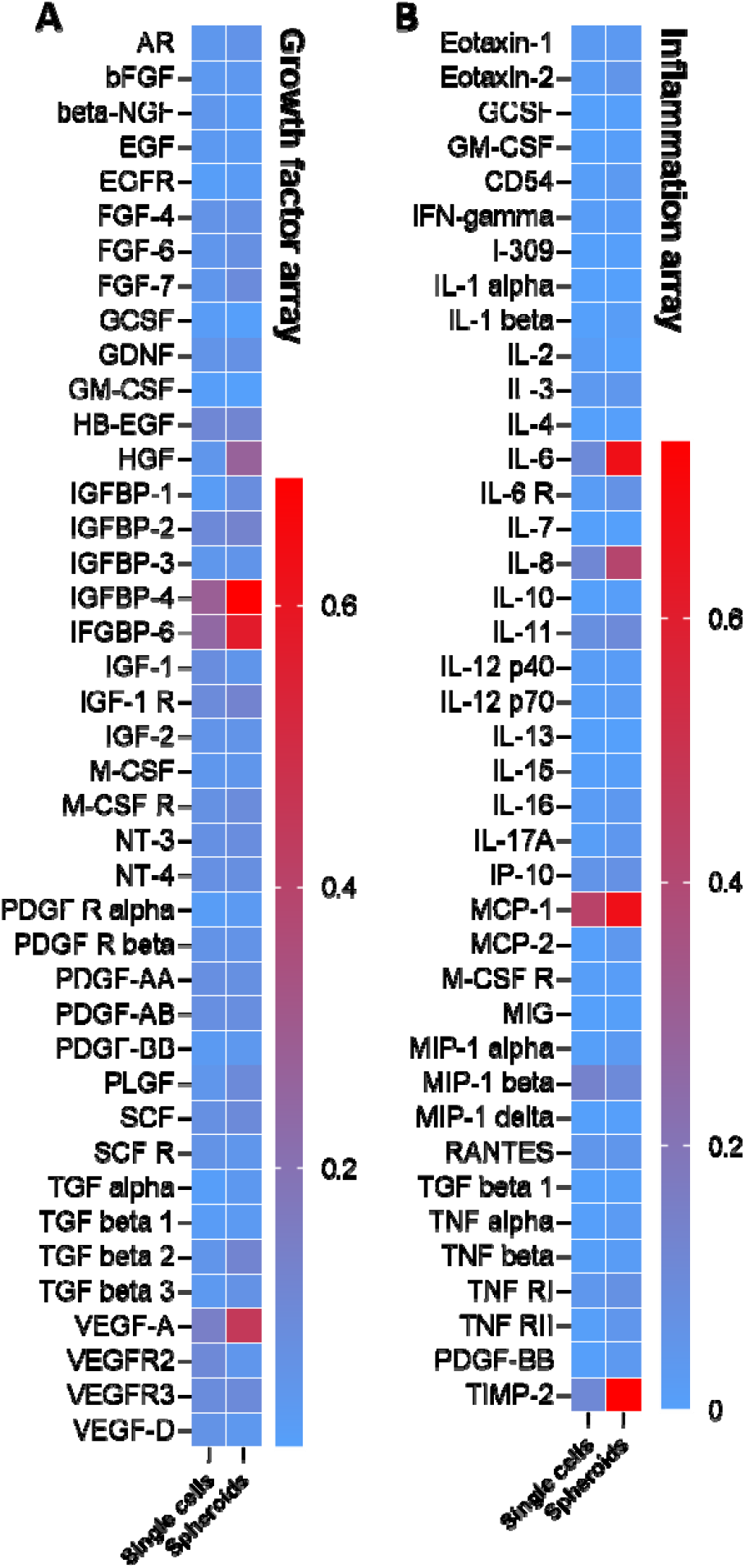
Growth factors and inflammation cytokines secreted in the CM prodiced from hMSCs encapsulated as single cells or assembled as spheroids in the macroporous hydrogels. (A) Heatmap of growth factors secretion. (B) Heatmap of inflammatory factors secrection. CM were collected from three hydrogel samples per condition.

The current study only explored MSC assembly and secretome profiles using the macroporous hydrogels templated by dissolvable PEGNB-Dopa microgels. Given that the PEGNB-Dopa can be readily crosslinked into hydrogels with any shapes or sizes through the efficient thiol-norbornene photopolymerization, we believe that this material platform has high potential in creating sophisticated internal structures via biofabrication techniques. For example, the dissolvable PEGNB-Dopa microgels can be encapsulated in sufficiently high density to create an interconnected porous structure. PEGNB-Dopa may also be photocrosslinked into other shapes (e.g., fibers, cylindrical posts, etc.) for serving as a rapidly dissolving sacrificial material compatible with *in situ* cell encapsulation. These hierarchical hydrogel structures may be used for basic science research or to facilitate tissue regeneration via enhancing cell infiltration or vascularization.

## 3. Conclusion

In conclusion, we have demonstrated a new biomaterial platform for rapid *in situ* formation of cell-laden macroporous hydrogels. Central to this platform was the use of PEGNB-Dopa microgels as the sacrificial porogens, which rapidly dissolved upon contacting with an aqueous solution. As the dissolution of the microgels did not rely on any external triggers or changes in environmental conditions (e.g., pH, temperature), this *in situ* pore-forming approach was high cytocompatible, as demonstrated by the *in situ* encapsulation of multiple cell types. Furthermore, adjusting the cell adhesiveness of the bulk hydrogels afforded the formation of solid cell spheroids or hollow spheres. The assembly of hMSC spheroids/spheres led to differential activation of the AKT pathway, with the solid spheroids exhibiting robust secretion of growth factors and certain cytokines. In summary, this platform provides an innovative method for forming cell-laden macroporous hydrogels for a variety of biomedical applications.

## Experimental Section

### Materials

PEG-OH (8-arm, 20 kDa, JenKem), carbic anhydride (Sigma-Aldrich), 5-norbornene-2-carboxylic acid (Sigma-Aldrich), 4-dimethylaminopyridine (DMAP, Alfa Aesar), dopamine hydrochloride (Sigma-Aldrich), tyramine hydrochloride (Chem-Impex), isopropylamine (TCI chemicals), N,N’-Diisopropylcarbodiimide (DIC, Chem-Impex), N,N’-Dicyclohexylcarbodiimide (DCC, Sigma-Aldrich), hydroxybenzotriazole (HOBt, ≥20 wt% water, Oakwood Chemical), N,N’-diisopropylethylamine (DIPEA, TCI chemicals), cysteine (Sigma-Aldrich), and lithium phenyl-2,4,6-trimethylbenzoylphosphinate (LAP, ≥95%, Sigma-Aldrich), 4-arms thiolated PEG (PEG4SH, 10 kDa, Laysan Bio), 5,5’-dithiobis-(2-nitrobenzoic acid) (Ellman’s reagent, Life Technologies), pyridine (Fisher), anhydrous tetrahydrofuran (THF, Fisher), anhydrous dimethylformamide (DMF, amine-free, Fisher), anhydrous dichloromethane (DCM, Fisher), methanol (Sigma-Aldrich) and diethyl ether (Fisher) were used as received.

### Synthesis of functionalized macromers

PEGNB was synthesized according to previously established protocols [39, 40]. Before conducting the reaction, PEG was distilled to remove excess water. To activate 5-norbornene-2-carboxylic acid (norbornene acid hereafter), 1.22 mL norbornene acid (10 mmol, 5 equiv.) was combined with 1.03 g DCC (5 mmol, 2.5 equiv.) in 30 mL anhydrous DCM under N_2_ for 1 h. Meanwhile, 5 g dry PEG-OH (hydroxy group 2 mmol), 122 mg DMAP (1 mmol, 0.5 equiv.), 810 µL pyridine (10 mmol, 5 equiv.) and 30 mL anhydrous DCM were charged in a separate round bottom flask. After 1 h, the white precipitate generated during the activation step was filtered out. The activated norbornene acid solution was then drop-added into PEG solution at 0 °C under N_2_ in dark. The reaction was proceeded overnight. To ensure high conjugation efficiency, another portion of activated norbornene acid was drop-added into the reaction the next day. After the reaction was completed, the solution was filtered and precipitated 2 times into diethyl ether. The crude product was vacuum dried and redissolved in pure water. The final product was harvested by dialyzing in pH 6 water for 3 days followed by lyophilization. The product was characterized by ^1^H NMR spectra.

PEGNB_CA_ and PEGNB-Dopa were synthesized using our published protocol [38]. Briefly, 10 g PEG-OH (hydroxyl group 0.4 mmol), 3.28 g carbic anhydride (2 mmol, 5 equiv.), 0.49 g DMAP (0.4 mmol, 1 equiv.), and 83 mL anhydrous THF were charged in a round-bottom flask equipped with a stir bar. The reaction was conducted at 60 °C. After 12 h, another portion of carbic anhydride and DMAP were added into the reaction. The reaction was continued for another 24 h. PEGNB_CA_ was retrieved via precipitation through diethyl ether twice and subsequently dried in vacuum. To synthesize PEGNB-Dopa, 1 g PEGNB_CA_ (carboxylic group 380 µmol), 148.8 µL DIC (950 µmol, 2.5 equiv.), 160 mg HOBt (950 µmol, 2.5 equiv.) and 10 mL amine-free DMF were charged into a reaction vial equipped with a stir bar. The activation of the acid group was blanketed under nitrogen gas for 2 h in dark. 180 mg dopamine hydrochloride (950 µmol, 2.5 equiv.), 165.4 µL DIPEA (950 µmol, 2.5 equiv.), and 23 mg DMAP (190 µmol, 0.5 equiv.) were dissolved in 1 mL amine-free DMF and then added into the reaction. The reaction was continued overnight. The product was dialyzed in methanol for one day and dried out in vacuum. The product was characterized by ^1^H NMR spectra (Bruker 500 MHz).

GelNB was synthesized according to our previous method.[49] Briefly, 2 g of type B gelatin was dissolved in PBS to make a 10 wt % gelatin solution. 0.6 g of carbic anhydride was added to the gelatin solution, and pH was adjusted to 7.5 – 8. The reaction was allowed to proceed for 24 h, after which the product was dialyzed against DI water for 3 days and lyophilized.

### Microfluidic chip fabrication

Microfluidic devices were designed and fabricated via conventional soft lithography as previously described.[58] Briefly, polydimethylsiloxane (PDMS) was poured onto a silicon wafer (Wafer World Inc, USA) with micro-structures designed in CAD, vacuumed for 60 minutes to remove entrapped air, then transferred to a 70 °C oven to cure overnight. PDMS replicas were then trimmed and punched with a 20 G dispensing needle (CML Supply, USA) to fashion inlets and outlets. After cleaning, PDMS replicas were bond to glass slides after exposing to oxygen plasma (Harrick Scientific, USA), then transferred to 70 °C oven to facilitate bonding. Channel dimensions (h × w) were 100 μm × 100 μm for cell and macromer stream, and 100 μm × 20 μm for the oil stream.

### Microgel fabrication

10 wt% PEGNB or PEGNB-Dopa, 20 mM PEG4SH, 7.2 mM LAP, with 0.1 wt% thiolated Rhodamine B was prepared as hydrogel-forming solution. PBS or cell-containing media was mixed with hydrogel-forming solution on-chip via a serpentine channel. Microgels with a final concentration of 5 wt% PEGNB or PEGNB-Dopa, 10 mM PEG4SH and 3.6 mM LAP were fabricated under constant flow rates for both PBS and macromer solution (2 μL/min) while varying oil phase flow rate to 4 μL/min, 12 μL/min, and 20 μL/min to create microgels of different sizes. Droplets were then exposed to 15 mW/cm^2^ UV for 20 seconds for gelation. Polymerized microgels were separated from oil and recovered into PBS by centrifugation on a 40 μm cell strainer.

### Macroporous or nanoporous hydrogel fabrication and characterization

After separating from oil, microgels were immediately encapsulated within a continuous hydrogel matrix to form macroporous hydrogels. Either PEGNB (5 wt%, 5 (R=0.5) or 10 mM (R=1) PEG4SH, 3.6 mM LAP), or GelNB (5 wt%, 0.5 (R=0.5) or 1 mM (R=1) PEG4SH, 3.6 mM LAP) was used to form contineous matrix, and 30 μl solution was exposed to 15 mW/cm^2^ UV light for 20 seconds in a 1 mL syringe with tip removed for gelation. Fluorescence images of the microgels were taken every 24 hours by using a fluorescent microscopy (Lionheart FX, BioTek) and fluorescence intensity was quantified in imageJ. Confocal microscopy was also utilized to verify the macroporous structure. For confocal imaging, both microgels and bulk gels were labeled with fluorescent dye (0.1 wt% thiolated-Rhodamine for microgels and 0.1 wt% FITC-PEG-SH for bulk gels). For measuring mechanical moduli, 45 μl hydrogel forming solution with/without microgels was injected in between two glass slides separated by 1 mm thick spacers. Hydrogels were swollen for 2 hours prior to measuring stiffness, which was done by using a digital rheometer (Bohlin CVO100, Malvern Instruments), with an 8 mm parallel plate geometry with a gap of 720 μm. For measuring swelling ratio, polymerized hydrogels were incubated in PBS for 2 hours before measuring wet weight. Dry weight of the hydrogels was measured after lyophilizing for 48 hours. Swelling ratio was calculated as the ratio between hydrogel wet weight and dry weight.

### Scanning electron microscopy (SEM) imaging

Non-degradable PEGNB bulk hydrogels and macroporous hydrogels were frozen in liquid nitrogen before lyophilization for 24 hours. And then the samples were imaged with a SEM after coating the hydrogel surface with gold.

### Cell culture, microencapsulation and encapsulation within macroporous hydrogels

Mouse MSCs were a generous gift from Dr. Hiroki Yokota (IUPUI BME). Human MSCs were isolated from deidentified bone marrow aspirate (Lonza). Mouse or human MSCs, lung epithelial cell line A549, pancreatic cancer cell lines Colo-357, and PANC-1 were cultured at 37 °C and 5% CO_2_ in Dulbecco’s Modified Eagle’s Medium (DMEM, HyClone) with low glucose (high glucose for Colo-357 and PANC-1) supplemented with 10 % fetal bovine serum (FBS, Corning, USA), 1% Antibiotic-Antimycotic (Life Technologies, USA). 2 ng/mL FGF-2 was used when culturing MSCs. Cell populations were subcultured every 3 days. To prepare cells for microencapsulation, cells were trypsinized, pelleted, and resuspended to a final concentration of 2×10^6^, 2×10^7^ and 4×10^7^ cells/mL in culture media or OptiPrep for microencapsulation as described above, except that cell stream was separated from macromer stream for homogeneous cell encapsulation. Flow rates for cell stream and macromer stream were kept at 2 μl/min, while the flow rate of oil stream was varied to control the diameter of microgels. The concentration of macromer solution was doubled to 10 wt% PEGNB-Dopa, 20 mM PEG4SH and 7.2 mM LAP, followed by encapsulation into macroporous hydrogels, where 30 μl solution was exposed to 15 mW/cm^2^ UV light for 20 seconds in a 1 mL syringe with tip cut off for gelation. Cell-laden macroporous hydrogels were then transferred and cultured in the incubator. To encapsulate MSCs into PEGNB or GelNB nanoporous hydrogels, 500,000 cells/mL were suspended into hydrogel-forming solutions for forming 5 wt% PEGNB or GelNB hydrogels with PEG4SH (R=0.5 or 1) as described above. 30 μl solution was exposed to 15 mW/cm^2^ UV light for 20 seconds in a 1 mL syringe with tip removed for gelation.

### Cell viability assay

Viability of encapsulated cells was measured by staining with Live/Dead viability assays (Biotium, USA), a cellular membrane integrity assay that stains live cells with green fluorescence and dead cells with red fluorescence. Cell viability was imaged using a fluorescence microscope and calculated based on three images.

### Immunofluorescence staining

Encapsulated cells were fixed in 4 % paraformaldehyde for one hour at room temperature on an orbital shaker, followed by washing with PBS three times. Then, cell-laden hydrogels were permeabilized in 0.5 % Triton X-100 at room temperature for 10 minutes. The hydrogels were then washed with PBS and stained with 100 nM Rhodamine Phalloidin (Cytoskeleton Inc) and DAPI (1:1000) for 3 hours at room temperature. After the final wash, the z-stack images were obtained with confocal microscopy. To stain E and N-cadherin, hydrogels were fixed, blocked (5 % BSA for 2 hours), permeabilized as described above, then incubated with mouse anti-E-cadherin (1:100) and rabbit anti-N-cadherin (1:100) primary antibodies in antibody diluent buffer at 4 °C overnight. After washing with 3 % BSA for 30 minutes at 4 °C, secondary antibodies Alexa 488 goat anti-mouse and Texas Red mouse anti-rabbit at a concentration of 10 μg/mL were applied overnight at 4 °C to visualize the presence of primary antibodies. After washing with PBS, hydrogels were imaged with confocal microscopy.

### AKT phosphorylation array

The AKT pathway phosphorylation array (RayBiotech US) was used to identify the phosphorylation level of proteins involved in the AKT signaling pathway, among structured hMSCs and hMSCs grow on 2D tissue culture plastics (TCPs) or 3D encapsulated as single cells in bulk PEGNB hydrogels. hMSCs were cultured for 7 days before lysis for the analysis of the AKT phosphorylation array. Cell lysate was detected with biotinylated detection antibodies and visualized via chemiluminescence. The intensity of the individual dot represented the amount of specific proteins and was normalized to the reference dots for quantification.

### Growth factor and inflammation arrays

Secretion of cytokines and inflammatory factors was measured from CM of hMSC spheroids and single cells encapsulated in bulk hydrogels. hMSCs were cultured in microporous or nanoporous hydrogels for 3 days before CM was collected. CM was detected with biotinylated detection antibodies and visualized via chemiluminescence. The intensity of the individual dot represented the amount of specific proteins and was normalized to the reference dots for quantification.

### Statistical analysis

All experiments were conducted independently for three times and results were reported are Mean ± SD. One-way ANOVA and unpaired t-test with Welch’s correction were used on Prism software to analyze the statistical significance of the data. *, **, ***, **** represent p<0.05, 0.01, 0.001, 0.0001, respectively.

## Acknowledgements

This work was supported in part by the National Cancer Institute (R01CA227737). The authors thank Dr. Hiroki Yokota for providing mouse MSCs.

